# Precision Genome Editing Unveils a Breakthrough in Reversing Antibiotic Resistance: CRISPR/Cas9 Targeting of Multi-Drug Resistance Genes in Methicillin-Resistant *Staphylococcus aureus*

**DOI:** 10.1101/2023.12.31.573511

**Authors:** Aysegul Ates, Cihan Tastan, Safak Ermertcan

**Affiliations:** Pharmeceutical Microbiology Department, Faculty of Pharmacy, Ege University, Izmir, Turkey; Transgenic Cell Technologies and Epigenetic Application and Research Center (TRGENMER), Üsküdar University, Istanbul, Turkey; Molecular Biology and Genetics Department, Faculty of Engineering and Natural Science, Üsküdar University, Istanbul, Turkey

**Keywords:** CRISPR-Cas9, Gene editing, Antibiotic resistance, Methicillin-resistant Staphylococcus aureus, CRISPR-Cas antimicrobials

## Abstract

Antibiotic resistance poses a global health crisis, limiting the efficacy of available therapeutic agents and hindering the development of new antibiotics. The pharmaceutical industry’s waning interest in antibiotic production further exacerbates this challenge. Addressing antibiotic resistance demands innovative solutions. Here, we explore the application of CRISPR-Cas-based antimicrobials as a pioneering approach to combat multidrug resistance. Our study focuses on the methicillin-resistant clinical Staphylococcus aureus (MRSA), a significant clinical threat. Utilizing CRISPR/Cas9 technology, we aimed to concurrently target methicillin (mecA), gentamicin (aacA), and ciprofloxacin (grlA, grlB) resistance genes, thereby altering the resistance profile and enhancing sensitivity to antibiotics. We engineered CRISPR plasmids containing sgRNAs specific to the target regions, which were subsequently electroporated into MRSA strains. Real-time Polymerase Chain Reaction (RT-PCR) assessed changes in resistance gene expression, while disk diffusion and broth microdilution methods determined alterations in resistance status. Western blotting analyzed changes in PBP2a expression, and Sanger sequence analysis confirmed mutations in target regions. Results revealed a statistically significant 1.5-fold decrease in mecA gene expression, 5.5-fold decrease in grlA gene, 6-fold decrease in grlB gene, and 4-fold decrease in aacA gene compared to the wild type strain, as determined by RT-PCR. Antibiotic susceptibility tests demonstrated the suppression of mecA, grlA, grlB, and aacA genes, resulting in the reversal of resistance to beta-lactam, quinolone, and aminoglycoside antibiotics. Western blot analysis showed 70% decrease in PBP2a expression, indicating the breakdown of beta-lactam resistance. Sanger sequence analysis confirmed point mutations in grlB and aacA genes, along with three-base mutations in grlA and mecA genes. Our findings underscore the potential of CRISPR/Cas9 technology to mitigate antibiotic resistance, offering a transformative strategy to restore the efficacy of existing antibiotics in the face of multidrug-resistant pathogens.

## Introduction

Antibiotic resistance, a formidable global challenge, transcends national borders and stands among the World Health Organization’s top 10 threats, presenting not only a health crisis but also a socio-economic dilemma due to the associated high mortality and morbidity rates.^1-3^ In 2019 alone, an alarming 5 million deaths resulted from untreatable infections owing to antibiotic resistance, underscoring the urgency of effective interventions.^3-4^ Projections indicate that by 2050, an annual toll of 10 million lives could be lost, threatening a 3.8% reduction in the gross domestic product, potentially impacting 24 million people if unchecked.^5^

While efforts to introduce novel antibiotics persist, the pharmaceutical industry faces formidable challenges, including the prolonged and costly process of drug development and the rapid emergence of resistance to newly introduced antimicrobials. Researchers navigating this landscape explore diverse strategies, such as drug repositioning and targeting metabolic or virulence pathways, to overcome these obstacles.^6-8^ Current antibiotics, while effective, lack the specificity to selectively eliminate targeted bacteria, disrupting not only pathogenic elements but also beneficial host microbiota.^9-12^ The global imperative for alternative, highly specific methodologies become evident in the face of these challenges.^13^

Advancements in genetic engineering technologies offer a promising avenue for the development of targeted antimicrobials against resistant microorganisms. The CRISPR/Cas system, initially characterized as bacteria’s adaptive immune system against phage genomes, emerged as a groundbreaking genome editing technology in 2012.^14-16^ Anticipated to become a novel defense mechanism in the battle against resistance, our study highlights the application of CRISPR antimicrobials in eliminating resistance to various antibiotics in diverse bacterial strains.

Motivated by the escalating global health crisis precipitated by antibiotic resistance and the diminishing therapeutic options, our investigation focuses on CRISPR antimicrobials as a potential solution. We specifically target Methicillin-Resistant Staphylococcus aureus (MRSA), identified by the World Health Organization as a critical pathogen necessitating urgent antibiotic development and research.^17^ With its genetic plasticity and virulence factors supporting pathogenesis, MRSA poses a severe threat, causing a spectrum of infections from uncomplicated to life-threatening.^18^ The crux of methicillin resistance lies in the presence of the PBP2a protein, encoded by the “mec” region on chromosomal DNA, which is absent in susceptible strains, rendering most MRSA strains resistant to a broad range of antibiotics.^19,20^

Our study specifically targets the methicillin resistance gene mecA, gentamicin resistance gene aacA, and ciprofloxacin resistance genes grlA and grlB in clinical MRSA isolates. The overarching goal is to sensitize these isolates to relevant antibiotics that have lost efficacy due to resistance, thus providing a promising avenue for addressing the challenges posed by multidrug-resistant pathogens.

## Materials and Methods

### Bacterial Strains, Plasmid & Primers

20 *S*.*aureus* strains isolated from clinical samples at the Department of Medical Microbiology of Ege University Hospital were included in this study. *S. aureus* ATCC 29213 was used as the control strain. All bacterial isolates were stored in Brain-Heart Infusion (BHI) broth medium with 10% glycerin at - 80°C until the study. pCasSA was a gift from Quanjiang Ji (Addgene plasmid # 98211). The primers used in the study are listed in Table S1.

### Identification of isolates and determination of antibiotic susceptibilities

Identification and antibiotic susceptibilities of clinical isolates were performed by Matrix assisted laser desorption ionisation time of flight massspectrometry (MALDITOF-MS) (Biomerieux, France) and VITEK-2 Compact (Biomerieux, France) automated system,respectively. VITEK-2 test results were confirmed by broth microdilution testing. *mec*A, *grl*A, *grl*B, *aac*A resistance genes in clinical isolates resistant to methicillin, ciprofloxacin, gentamicin were detected by conventional PCR. A representative clinical MRSA isolate was selected and used in study.

### Restrictional Digestion

To reduce the risk of false positive results in the ligation of gRNAs into the plasmid, the plasmid was linearised with *Bsa*I (NEB,UK). Linearize product was visualized by electrophoresis in 1% agarose gels and then purified from agarose gel. Purification was performed using the Plus Gel Eluted Kit (GMbiolab, Taiwan).

### Phosphorylation, annealing of oligos and plasmid ligation

On-target and off-target scores were considered and oligos specific to each target site were designed from Benchling.com. These oligos were synthesized with GAAA insertion at the 5’ end of the first sequence and CAAA extension at the 5’ end of the second chain to bind to the sticky ends formed in the plasmid after restriction digestion (Table S2). To reduce the possibility of false positives, instead of direct Golden gate cloning, oligos was inserted by ligation into linearize plasmid. The ligation product was transferred into *E*.*coli* DH10B chemically competent cells by heat shock transformation. After transformation, cells were inoculated on LB agar containing 50 μg/mL kanamycin. After incubated at 37°C, sample colonies were selected on LB agar containing 50 μg/mL kanamycin for plasmid isolation. Isolation was performed using Monarch Plasmid Miniprep Kit (NEB, UK).

### Colony PCR

After plasmid isolation, colony PCR was used to confirm whether the targeted ligation occurred in the isolated plasmids. For this purpose, colony PCR was performed to amplify a 120 bp region with appropriate primers, with the Forward primer being the site of plasmid ligation.

### Preparation of electrocompetent Methicillin Resistant *S. aureus* cells

A single colony of MRSA was inoculated into TSB medium. After 18 hours of incubation, an appropriate volume of the broth culture was taken and diluted 1/100 in TSB medium and incubated in shaking incubator at 30 °C for 2-3 hours (i.e. OD600: ∼ 0.4). When the targeted OD600 value was reached, the culture was incubated dry ice for 10 minutes. The culture was centrifuged at 5000 rpm for 5 minutes. The supernatant was removed and cells were suspended in 500 mM sucrose solution. After centrifugation and re-suspension steps were repeated twice, the supernatant was discarded and the pellets were suspended in 0.5 M sucrose, aliquoted and stored at -80°C until electroporation.^21^

### Electroporation

The electrocompetent MRSA suspension was mixed with pCasSA (∼2 μg) containing the insert unit appropriate for the target genes by gently tapping the tube. The mixture was transferred to a 1 mm electroporation cuvette at room temperature. The cells in the cuvette were pulsed using Eporator (EPPENDORF-Eporator, Germany) at 21 kV/cm, 5.0 ms. After electroporation, the cells were incubated in TSB medium at 30°C for 1.5 hours. The transformant cell culture was then inoculated with a drigalski spatula into TSA medium containing 5 μg/mL chloramphenicol. The plates were incubated at 30°C for 48 hours.^21^

### Investigation of Expression Levels of Resistance Genes by Quantitative Real-Time Reverse Transcriptase PCR (RT-qPCR)

RNA isolation from Wild Type (WT) clinical MRSA isolate and Knock-out (KO) strain was performed using the Total RNA purification kit (GMbiolab, Taiwan). Concentrations and purities of RNA isolates were measured on a Beckman Coulter DU-730 (USA). The cDNA was synthesized from RNA at the appropriate concentration using Onescript Plus Reverse Transcriptase cDNA Synthesis Kit (ABM, USA) according to the manufacturer recommendations. The cDNAs synthesized from total RNAs were used for RT-qPCR with 2X Magic SYBR Mix (Procomcure, Australia) kit. cDNA concentrations were measured, equalized and then, added to a 96-well Armadillo (Thermo Scientific, USA) microplate. the master mix was added and the plate was placed in a LightCycler 480 II (Roche). The results were evaluated by Delta delta CT method considering the Cp values obtained from LightCycler 480 Software release 1.5.0. SP3. The primers for *mec*A, *aac*A, *grl*A, *grl*B and housekeeping gene 16srRNA utilized in qPCR experiments are shown in Table S1. Using 16S rRNA as housekeeping gene, changes in the expression levels of antibiotic resistance genes were statistically evaluated by normalization with ΔΔCT method.

### Phenotypic antibiotic susceptibility tests

The effect of genomic modification by CRISPR/Cas technology on the resistance profile was examined by phenotypic antibiotic susceptibility tests. Broth microdilution and disk diffusion methods for WT and KO isolates were performed according to EUCAST guidelines. *S. aureus* ATCC 29213 was used as control strain.

### Broth microdilution method

The broth microdilution method was performed in accordance with EUCAST and ISO (20776) criteria. The turbidities of WT and KO strains were adjusted to McFarland 0.5 with 0.9% saline. Bacterial inoculums were diluted 1/100 with MHB to a final concentration of 5x10^5^ CFU/mL. Ciprofloxacin, oxacillin, gentamicin solutions with final concentrations (2048 μg/ml) were prepared with sterile distilled water. MHB was added to all wells of a U-bottom 96-well sterile microplate and then oxacillin, gentamicin and ciprofloxacin solutions were added only to the first wells of the microplate. Serial dilution was performed. Bacterial suspensions were inoculated into the wells. Sterility control and growth controls were included. After incubated at 37°C, 24-48 h, the lowest concentration without growth was determined as the minimum inhibitory concentration (MIC-μg/ml).^22,23^

### Disk diffusion method

WT and KO strains were adjusted to a standard turbidity of McFarland 0.5 with 0.9% saline. The strains were inoculated onto MHA medium with swab. After the media were allowed to dry for a few minutes, cefoxitin (30 μg-FOX-30), gentamicin (10 μg-CN-10) and norfloxacin (10 μg-NOR-10) disks were placed on the media using sterile forceps. After incubation at 37 °C for 24-48 hours, the zone diameters around the disks were measured and evaluated.^24^

### Protein purification and Western Blot

Western blotting was performed to evaluate the expression change in PBP2a protein in KO strains. For this purpose, protein isolation was performed from WT and KO strains.^25^ Bradford Commasie Blue method was used to determine protein concentrations. Pierce™ Coomassie (Bradford) Protein Assay Kit (Thermo Scientific, USA) was applied according to the manufacturer recommendations. Protein concentrations were calculated using absorbance values. Proteins were brought to equal concentrations before loading (11 μg). Protein molecular weight marker (GenemarkBio, Taiwan) was added to the first well, WT protein and KO protein mixture were added to the other wells in triplicate. Electrophoresis was performed with Bio-Rad Protean-II electrophoresis system at 80 V for 10 minutes, then 100 V for 120 minutes. After SDS-PAGE was completed, proteins were transferred to PVDF (ThermoScientific, USA) membranes (+4 °C, 100 V, 2h). Non-specific binding was prevented after transfer. Then, the primary antibody (Mouse Anti-Pbp2a, Raybiotech, USA) was diluted 1/1000 in 5% milk powder and added to the membrane. The membrane was then incubated overnight at +4°C on an orbital shaker. At the end of the incubation, the membrane was washed 3 times with 1xTBS-T for 10 min incubations. Secondary antibody (Anti-mouse IgG, Jackson ImmunoResearch, UK) diluted 1/10000 was added to the membrane and incubated on a shaker at room temperature for 2 hours. At the end of incubation, the membrane was washed 3 times with 1XTBS-T at 10 min intervals. Luminol/Enhancer from the chemiluminescence substrate kit (SuperSignal West Pico Plus, Thermo Scientific, USA) and Peroxidase Buffer were added to the membrane in a 1:1 ratio and kept on the shaker for 5 minutes. Imaging was performed with Fusion FX-7 (Vilbert Lourmat, France).

### Sanger sequence analysis

Genomic DNA isolation of KO *S. aureus* strains was performed using Nucleospin Microbial DNA kit (Macherey-Nagel, Germany). In order to examine the mutation caused by the CRISPR/Cas system in the target gene regions, regions of approximately 1000bp were selected to include each gene region. The relevant regions were amplified by PCR using primers designed for this study and shown in Table S1 Q5 High Fidelity DNA polymerase (NEB) was used for sequence analysis with appropriate purity and accuracy.

### Statistical Analysis

All experiments were made in triplicates. standard deviations (SDs) are calculated by GraphPad Prism 5 software (San Diego, CA, USA). Statistical calculates were analyzed by one-way ANOVA tests. p<0.5 was accepted as statistically significant.

## Results

### Quantitative Real-Time Reverse Transcriptase PCR (RT-qPCR)

To quantitatively analyze changes in gene expression, we employed Real-Time Reverse Transcriptase PCR (RT-qPCR) using 16S rRNA as a housekeeping gene. The expression levels of targeted genes were statistically evaluated by normalizing with the ΔΔCT method, providing a robust framework for accurate comparisons. Figure 1 illustrates the gene expression changes in Knockout (KO) strains in comparison to Wild Type (WT) strains.

**FIG 1.**
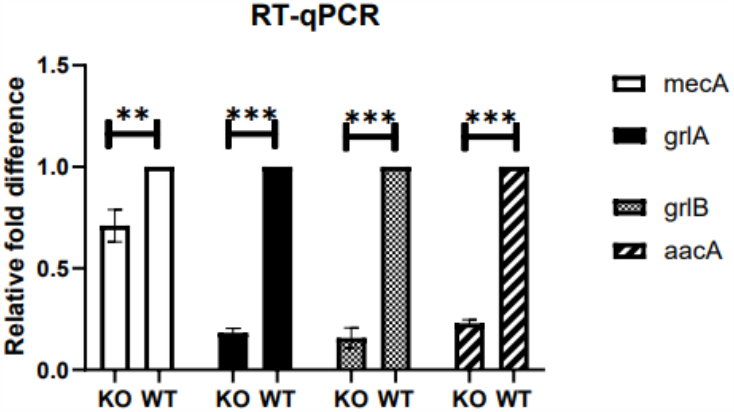
Changes in the expression ratio of related genes in RT-PCR (KO: Knock-out, WT: Wild type, *: p < 0.01, ***: p < 0.001)

In the KO strain, a discernible impact was observed on the expression levels of key resistance genes. Relative to the WT isolate, a significant 1.5-fold decrease in mecA gene expression was noted, indicative of the successful targeting and suppression of methicillin resistance. Moreover, the KO strain exhibited a remarkable 5.5-fold reduction in grlA gene expression, a 6-fold decrease in grlB gene expression, and a 4-fold decrease in aacA gene expression compared to the WT strain.

These findings underscore the precision and efficacy of the CRISPR/Cas9-mediated genome editing approach in selectively modulating the expression of resistance-associated genes. The observed reductions in gene expression levels highlight the successful alteration of the resistance profile in the KO strains, providing crucial insights into the potential of CRISPR technology as a tool for combating antibiotic resistance.

### Phenotypic antibiotic susceptibility tests

#### Broth microdilution and disk diffusion method

The results of broth microdilution and disk diffusion tests for clinical strains and quality control strains were evaluated in Table 1 in accordance with EUCAST guidelines.

**Table 1.** Antibiotic susceptibility test results.

| STRAINS<br>→ | EUCAST STANDARDS |  |  |  | TEST RESULTS |  |  |  |  |  |
| --- | --- | --- | --- | --- | --- | --- | --- | --- | --- | --- |
|  | *S. aureus*<br>ATCC 29213 |  | *S. aureus* |  | *S. aureus*<br>ATCC 29213 |  | *S. aureus*<br>(WT) |  | *S. aureus*<br>(KO) |  |
| ↓<br>ANTIBIOTICS | MIC<br>(mg/L) | Zone<br>diameter<br>(mm) | MIC<br>(mg/L) | Zone<br>diameter<br>(mm) | MIC<br>(mg/L) | Zone<br>diameter<br>(mm) | MIC<br>(mg/L) | Zone<br>diameter<br>(mm) | MIC<br>(mg/L) | Zone<br>diameter<br>(mm) |
| OXACILLIN | 0.125-0.5 | - | >2:R | - | 0.5 | - | 128 | - | 1 | - |
| CEFOXITIN | - | 24-30 | - | ≥22:S<br><22:R | - | 28 | - | 6 | - | 28 |
| CIPROFLOXACIN | 0.125-0.5 | - | ≤0.001:<br>S>1:R | ≥17:S<br><17:R | 0.5 | - | >512 | - | 1 | - |
| NORFLOXACIN | - | 18-24 | - | ≥17:S<br><17:R | - | 19 | - | 6 | - | 30 |
| GENTAMICIN | 0.125-1 | 19-25 | ≤2:S<br>>2:R | ≥18:S<br><18:R | 0.5 | 24 | >512 | 6 | 4 | 22 |

Antibiotic susceptibility tests unveiled a remarkable transformation in the resistance profile of the Wild Type (WT) strain, which exhibited resistance to all antibiotics tested. In contrast, the Knockout (KO) strain, engineered through CRISPR/Cas technology, demonstrated a notable shift towards susceptibility across various antibiotics. Notably, the cefoxitin zone diameter witnessed a substantial increase, coupled with a 128-fold decrease in oxacillin MIC values in the KO strain, signifying the elimination of methicillin resistance. The precision of CRISPR/Cas technology in disrupting resistance mechanisms was further evidenced by a significant decrease in ciprofloxacin MIC values and an augmented zone diameter for norfloxacin, resulting in the reclassification of bacteria as susceptible, effectively abolishing ciprofloxacin resistance. Furthermore, aminoglycoside resistance was decisively broken in the KO strain, as evident from a notable increase in the gentamicin zone diameter and an approximately 512-fold reduction in MIC values. The observed phenotypic changes were directly attributed to the targeted suppression of mecA, grlA, grlB, and aacA genes, collectively leading to the cessation of methicillin, quinolone, and aminoglycoside resistance. These results conclusively demonstrate the efficacy of CRISPR/Cas9-mediated genetic intervention in effecting a paradigm shift from resistance to susceptibility in response to relevant antibiotics. The successful elimination of multiple drug resistances underscores the potential of this innovative approach in addressing the formidable challenge of antibiotic resistance.

#### Western Blotting

The CRISPR modification targeting the mecA gene yielded profound effects on both gene expression and phenotypic resistance, as demonstrated through a comprehensive set of analyses. Real-Time Polymerase Chain Reaction (RT-PCR) results revealed a significant reduction in mecA gene expression, indicative of the successful modulation of methicillin resistance at the molecular level. Subsequent phenotypic antibiotic susceptibility tests confirmed the breakthrough, highlighting the abolition of methicillin resistance in the Knockout (KO) strains.

To further elucidate the impact of mecA gene modification, Western blot analysis was conducted to assess the expression of Penicillin-Binding Protein 2a (PBP2a). Figure 2 illustrates a conspicuous decrease in PBP2a expression in the KO-mecA strain, underscoring the efficacy of the CRISPR-mediated genetic intervention. Image analysis using ImageJ 8 software and normalization according to Spa allowed for a precise quantification of the expression changes.

**FIG 2.**
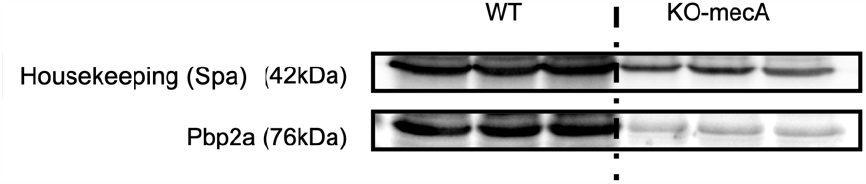
Western Blot membrane image (WT: Wild Type, KO-mecA: mecA gene suppressed strain)

As depicted in Figure 3, a striking 70% reduction in PBP2a expression was observed in the KO strain compared to the Wild Type (WT), further validating the successful disruption of methicillin resistance.

**FIG 3.**
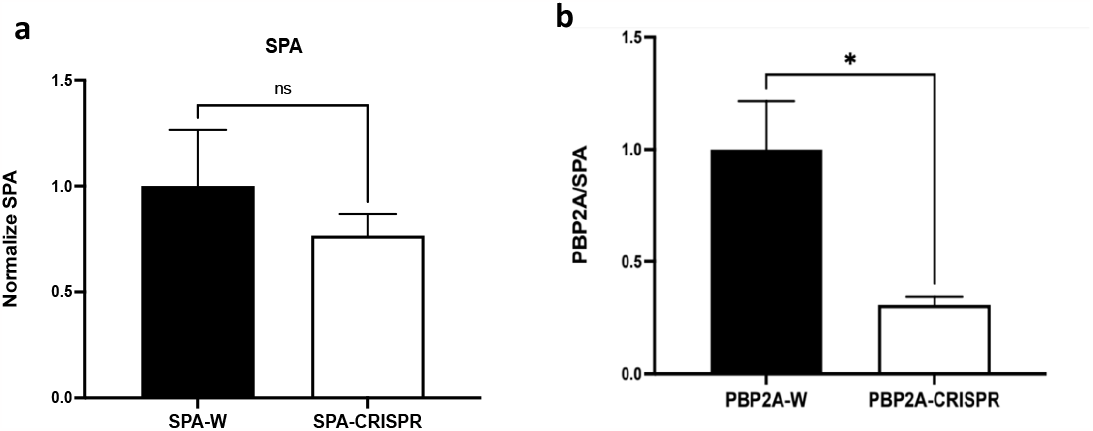
**a**.Spa, **b**.PBP2a expression ratio changes (SPA-W: Spa in wild type, Spa-CRISPR: Spa in mecA repressed strain, PBP2A-W: Pbp2a in wild type strain, PBP2A-CRISPR: Pbp2a in mecA gene suppressed strain, ns:Not statistically significant, not significant, *: Statistically significant (p < .05))

These findings not only showcase the potency of CRISPR technology in achieving targeted genetic modifications but also emphasize its transformative potential in addressing antibiotic resistance at both the genetic and phenotypic levels. The ability to modulate the expression of key resistance genes holds promise for advancing precision therapies against multidrug-resistant pathogens.

#### Sanger sequence analysis

The results of Sanger sequence analysis were compared with the sequences in the NCBI gene bank (Accession number: BA000017, AP003367.1, D67075.1) and BLAST analysis was performed. The mutations obtained are shown in Figure 4.

**FIG 4.**
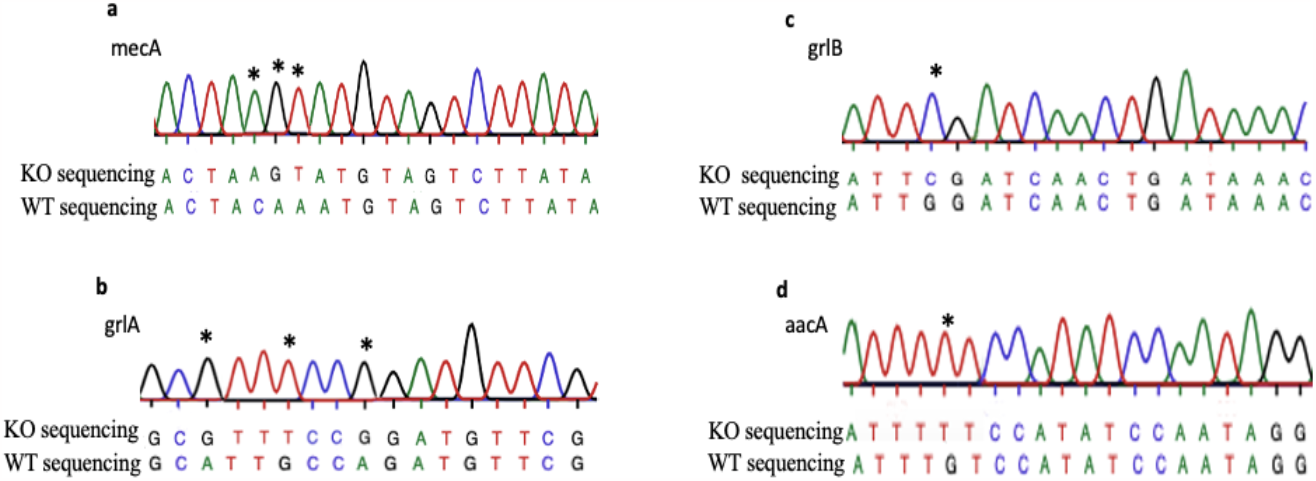
Mutations detected in KO isolate. **(a)** In the KO strain, positions 33., 34. and 35. of the mecA gene occurred C→A, A→G, A→T point mutations, respectively. **(b)** In the KO strain, positions 90, 93 and 96 of the grlA gene occurred A→G, G→T ve A→G point mutations, respectively. **(c, d)** KO strain, positions 88 of the grlB gene and positions 220 of the aacA gene appeared G→C, G→T point mutations, respectively.

## Discussion

Antibiotic resistance remains a critical global health and socio-economic concern, with a growing urgency to address the imminent challenges it poses. The stagnation in new antimicrobial development, despite the escalating rates of antibiotic resistance, underscores the pressing need for innovative therapeutic strategies to mitigate the impending crisis. Within the realm of antibiotic resistance, Methicillin-Resistant Staphylococcus aureus (MRSA) infections, characterized by both virulence and drug resistance, demand the exploration of groundbreaking treatment methods. CRISPR/Cas technology emerges as a pivotal component among innovative approaches to combat antibiotic resistance, asserting its influence in diverse scientific domains. As a versatile tool in molecular biology, CRISPR technology plays a pivotal role in microbiology, shaping the trajectory of future interventions against antibiotic resistance. The current study specifically focuses on the mecA gene, a key determinant of beta-lactam antibiotic resistance, emphasizing the need for innovative treatment strategies for MRSA infections.

Our investigation aligns with the pioneering work of Bikard et al. (2014), who targeted mecA and aph-3 genes using phagmids and the CRISPR/Cas system. While differences exist between our study and theirs, both endeavors share a common objective of suppressing the mecA resistance gene.^26^ In our study, a 30% suppression rate was achieved, leading to a substantial decrease in gene expression and consequent phenotypic shifts in antibiotic susceptibility. Wang and Nicholaou (2017) similarly aimed to repress the mecA gene, demonstrating a 77% decrease in gene expression.^27^ Our findings complement this work, showing a 30% reduction in mecA expression. Notably, our study extends beyond gene expression analysis to encompass phenotypic shifts, revealing a significant decrease in MIC values and expanded antibiotic zones in disk diffusion assays, indicative of a transformative impact on antibiotic resistance.

In contrast, Guan et al. (2017) achieved a cytotoxic effect on bacteria through the chromosomal targeting of the mecA gene.^28^ Our study, while distinct, resonates with their objective of disrupting oxacillin resistance, resulting in susceptibility classification. The differences between the studies highlight the various avenues through which CRISPR technology can be employed to achieve desired outcomes. The study by Kang et al. (2017) introduced a novel CRISPR nanoparticle (Cr-Nanocomplex) for targeting the mecA gene, showcasing its enhanced activity compared to the conventional CRISPR/Cas system.^29^ While our study aligns with the goal of targeting mecA, the focus extends to phenotypic shifts, emphasizing a significant reduction in MIC values. The diverse methodologies employed in CRISPR-based studies highlight the adaptability of this technology for different applications. Kiga et al. (2020) employed the Cas13 protein, a type VI, class 2 enzyme, for sequence-specific bactericidal effects.^30^ While their approach differs, our study complements their findings by showcasing the CRISPR/Cas9 technology’s efficacy in suppressing multiple resistance genes and achieving phenotypic changes in antibiotic susceptibility. Comparatively, our study distinguishes itself by concurrently targeting mecA, grlA, grlB, and aacA resistance genes. The observed reductions in gene expression and the subsequent phenotypic shifts, particularly in cefoxitin, ciprofloxacin, and gentamicin resistance, underscore the versatility and effectiveness of CRISPR technology against multiple drug resistances.

In conclusion, this study signifies a pioneering effort in the application of CRISPR/Cas9 technology to simultaneously suppress multiple resistance genes in clinical MRSA isolates. The demonstrated success in eliminating methicillin, ciprofloxacin, and gentamicin resistance, coupled with the statistically significant reductions in gene and protein expression, provides compelling evidence for the transformative potential of CRISPR-based approaches in the ongoing battle against antibiotic resistance. The findings contribute valuable insights into the development of precision therapies and underscore CRISPR technology as a promising therapeutic avenue for combating antibiotic resistance.

## Conclusions

The quest for innovative therapeutics to address the escalating challenge of antibiotic resistance has propelled extensive research into CRISPR/Cas antimicrobials. Our study, aligned with various investigations in the literature, underscores the transformative potential of CRISPR/Cas technology in targeting resistance genes within microbial populations, effectively transitioning them from resistance to susceptibility. Described as “DNA surgery at the nano-level,” CRISPR technology stands out for its facile application, cost-effectiveness, and programmable nature, offering specific modifications within targeted gene regions. As we envision the future trajectory of CRISPR antimicrobials, the prospect of simultaneously targeting multiple pathogens in a sequence-specific manner emerges as a promising avenue. This breakthrough technology holds the potential to revolutionize treatment protocols, allowing the simultaneous suppression of more than one gene within a bacterial species. Such advancements are anticipated to usher in groundbreaking developments in the realm of infectious disease treatment.

Our study contributes foundational data that lays the groundwork for future investigations. The path forward may involve exploring the use of CRISPR antimicrobials in combination with traditional antibiotics, overcoming limitations imposed by resistance development. Additionally, investigations into phage cocktails containing CRISPR/Cas antimicrobials and their synergistic effects with nanopharmaceuticals present exciting avenues for exploration. As we peer into the near future, the realm of CRISPR/Cas antimicrobials promises to inspire and catalyze exciting projects. The potential to reshape treatment paradigms and confront antibiotic resistance head-on positions CRISPR technology as a transformative force in the ongoing battle against infectious diseases. The convergence of CRISPR/Cas antimicrobials with other therapeutic modalities holds the key to unlocking novel strategies and ushering in a new era of precision therapeutics.

## Acknowledgments

pCasSA was a gift from Quanjiang Ji (Addgene plasmid # 98211). We are grateful to Associate Professor Quanjiang Ji.

## Authors’ Contributions

A.A., C.T., and Ş.E. designed the experiments. A.A. performed microbiology and molecular biology experiments. A.A., C.T., and Ş.E. analyzed the data and wrote the manuscript. All authors read and approved the final version of the manuscript

## Author Disclosure Statement

The authors declare no competing interests.

## Funding statement

Thss research was supported from the Ege Unsversity Scientific Research Projects Coordination by Grant number TDK-2020-21898

## Notes

### Competing Interest Statement

The authors have declared no competing interest.

## References

1- Katz L, Baltz RH. Natural product discovery: past, present, and future. J Ind Microbiol Biotechnol 2016;43(2-3): 155–176. doi: 10.1007/s10295-015-1723-5

2- Prescott JF. The resistance tsunami, antimicrobial stewardship, and the golden age of microbiology.Vet Microbiol 2014;171(3-4):273–278. doi:10.1016/j.vetmic.2014.02.035

3- Aslam B, Wang W, Arshad MI, et al. (2018). Antibiotic resistance: a rundown of a global crisis. Infect Drug Resist 2018;11:1645–1658. doi:10.2147/IDR.S173867

4- 10 global health issues to track in 2021. World Health Organization. [Online]. https://www.who.int/news-room/spotlight/10-global-health-issues-to-track-in-2021

5- O’Neill J. Tackling Drug-Resistant Infections Globally: Final Report and Recommendations, The Review on Antimicrobial Resistance. London, UK: World Health Organization, 2016

6- Su Z, Honek JF. Emerging bacterial enzyme targets. Cur opin investig 2007;8(2),140–149.

7- World Health Organization (WHO). Antimicrobial Resistance: Global Report on Surveillance. Switzerland: World Health Organization,2014

8- World Health Organization (WHO). 2015. Global Action Plan on Antimicrobial Resistance. Switzerland: World Health Organization,2014

9- Kohanski MA, Dwyer DJ, & Collins JJ. How antibiotics kill bacteria: from targets to networks. Nature Rev Microbiol 2010;8(6):423–435. doi:10.1038/nrmicro2333

10- Lock RL, Harry EJ. Cell-division inhibitors: new insights for future antibiotics. Nat Rev Drug Discov 2008;7(4):324–338. doi:10.1038/nrd2510

11- Njoroge J, Sperandio V. Jamming bacterial communication: new approaches for the treatment of infectious diseases. EMBO Mol Med 2009;1(4):201–210. doi:10.1002/emmm.200900032

12- Schimmel P, Tao J, Hill J. Aminoacyl tRNA synthetases as targets for new anti-infectives. FASEB J 1998;12(15), 1599–1609.

13- Duan C, Cao H, Zhang LH, Xu Z. Harnessing the CRISPR-Cas Systems to Combat Antimicrobial Resistance. Front Microbiol 2021;12, 716064. doi:10.3389/fmicb.2021.716064

14- Riordan SM, Heruth DP, Zhang, LQ, Ye, SQ. Application of CRISPR/Cas9 for biomedical discoveries. Cell Biosci 2015;5(1), 1–11. doi:10.1186/s13578-015-0027-9

15- Wilkinson R, Wiedenheft B. A CRISPR method for genome engineering. F1000Prime Reports 2014;6:3. doi:10.12703/P6-3

16- Jinek M, Chylinski K, Fonfara I. A programmable dual-RNA-guided DNA endonuclease in adaptive bacterial immunity. Science 2012;337(609):816–821. doi:10.1126/science.1225829

17- WHO publishes list of bacteria for which new antibsotics are urgently needed. World Health Organization. [Online]. https://www.who.int/news/item/27-02-2017-who-publishes-list-of-bacteria-for-which-new-antibiotics-are-urgently-needed

18- Moore PC, Lindsay JA. Genetic variation among hospital isolates of methicillin-sensitive Staphylococcus aureus: evidence for horizontal transfer of virulence genes. J Clin Microbiol 2001; 39(8):2760–2767. doi:10.1128/JCM.39.8.2760-2767.2001

19- Lade H, Joo HS, Kim JS. (2022). Molecular Basis of Non-β-Lactam Antibiotics Resistance in Staphylococcus aureus.Antibiotics (Basel) 2022;11(10):1378. doi:10.3390/antibiotics11101378

20- Sader, H. S., Watters, A. A. Fritsche, et al. Penicillin-binding protein 2a of methicillin-resistant Staphylococcus aureus. IUBMB life 2014;66(8):572–577. doi:10.1002/iub.1289

21- Chen W, Zhang Y, Yeo WS, Bae T, Ji Q. Rapid and Efficient Genome Editing in Staphylococcus aureus by Using an Engineered CRISPR/Cas9 System. J Am Chem Soc 2017;139(10):3790–3795. Doi:10.1021/jacs.6b13317

22- ISO 20776-1:2019 .Online]. https://standards.steh.as/catalog/standards/ssst/cfc459ae-0dee-4fb2-8ed7-e9458f6c7d6b/iso-20776-1-2019

23- The European Committee on Antimicrobsal Susceptibility Testing. Breakpoint tables for interpretation of MICs and zone diameters. Version 13.0, 2023. [Online]. http://www.eucast.org

24- The EUCAST Disk Diffusion Method for Antimicrobsal Susceptibility Testing Version 11.0 (January 2023). [Online]. http://www.eucast.org

25- Öztürk İ, Eraç Y, Ballar Kırmızibayrak P, Ermertcan Ş. Nonsteroidal antiinflammatory drugs alter antibiotic susceptibility and expression of virulence related genes and protein A of Staphylococcus aureus. Turk J Med Sci 2021;51(2):835–847. doi:10.3906/sag-2003-60

26- Bskard D, Eular CW, Jiang W, et al. Bikard, D., Euler, C. W., Jiang, W., Nussenzweig, P. M., Goldberg, G. W., Duportet, X.,Fischetti, V. A., & Marraffins, L. A. Exploiting CRISPR-Cas nucleases to produce sequence-specific antimicrobials. Nat Biotechnol 2014;32(11):1146–1150. dos:10.1038/nbt.3043

27- Wang K, Nicholaou M. Suppression of Antimicrobial Resistance in MRSA Using CRISPR-dCas9. Clin Lab Sci 2017;30(4),207–213. doi:10.29074/ascls.30.4.207

28- Guan J, Wang W, Sun B. Chromosomal Targeting by the Type III-A CRISPR-Cas System Can Reshape Genomes in Staphylococcus aureus. mSphere 2017;2(6):e00403–17. doi:10.1128/mSphere.00403-17

29- Kang YK, Kwon K, Ryu JS, et al. Nonviral Genome Editing Based on a Polymer-Derivatized CRISPR Nanocomplex for Targeting Bacterial Pathogens and Antibiotic Ressitance. Bioconjug Chem 2018;29(11):3936. doi: 10.1021/acs.bioconjchem.6b00676

30- Ksga K, Tan XE, Ibarra-Chávez R, et al. Development of CRISPR-Cas13a-based antimicrobials capable of sequence-specific killing of target bacteria. Nat Commun 2020;11(1):2934. doi:10.1038/s41467-020-16731-6

