## Supplemental Table for "Precision Genome Editing Unveils a Breakthrough in Reversing Antibiotic Resistance: CRISPR/Cas9 Targeting of Multi-Drug Resistance Genes in Methicillin-Resistant *Staphylococcus aureus*"

**Table S1. Primers used in this study.**

|  | **Primer** | **Sequence**  **(5’- 3’)** | **Product size**  **(bp)** |
| --- | --- | --- | --- |
| **Colony PCR** | ***grl*A-F** | GAAAATTGGATCAACTGATAAACG | 120 |
|  | ***grl*B-F** | GAAAGCATTGCCAGATGTTCGTGA | 120 |
|  | ***aac*A-F** | GAAATATATTTGTCCATATCCAAT | 120 |
|  | ***mec*A-F** | GAAAACTACAACTATTAAAATAAG | 120 |
|  | **PCR-R** | GGGTATGGACAGATCTCAAAAAAAGCAC | 120 |
| **Sanger Sequencing** | ***grl*B-F** | TGATACTTGCATTTTACGCTGAT | 1066 |
|  | ***grl*B-R** | ACAACAGCTGTTAAACCTTCACG |  |
|  | ***grl*A-F** | TGTTGAGTTTGGTATGCAAGAGG | 1167 |
|  | ***grl*A-R** | AACAACCTCAATTTGGTGATTCA |  |
|  | ***aac*A-F** | CACAGGAGTCTGGACTTGACTCA | 1188 |
|  | ***aac*A-R** | GCCCTTATTGCTCTTGGATTA |  |
|  | ***mec*A-F** | CGTTACAGTGTCACTTTCAACAT | 1146 |
|  | ***mec*A-R** | AACGATTGTGACACGATAGCC |  |
| **Housekeeping gene** | **16s*rRNA*-F** | ACGTGGATAACCTACCTATAAGACTGGGAT | 94 |
|  | **16s*rRNA*** | TACCTTACCAACTAGCTAATGCAGCG |  |

**Table S2. Oligos used in this study**

| **Name** | **Sequence**  **(5’ – 3’)** | **Description** |
| --- | --- | --- |
| ***grl*BF** | **GAAA**ATTGGATCAACTGATAAACG | grlB spacer for gene deletion |
| ***grl*BR** | **AAAC**CGTTTATCAGTTGATCCAAT | grlB spacer for gene deletion |
| ***grl*AF** | **GAAA**GCATTGCCAGATGTTCGTGA | grlA spacer for gene deletion |
| ***grl*AR** | **AAAC**TCACGAACATCTGGCAATGC | grlA spacer for gene deletion |
| ***aac*AF** | **GAAA**AGAGTATTAGAATTTTATGG | aacA spacer for gene deletion |
| ***aac*AR** | **AAAC**CCATAAAATTCTAATACTCT | aacA spacer for gene deletion |
| ***mec*AF** | **GAAA**GAAAATTTAAAATCAGAACG | mecA spacer for gene deletion |
| ***mec*AR** | **AAAC**CGTTCTGATTTTAAATTTTC | mecA spacer for gene deletion |
